# Low-frequency direct cortical stimulation of left superior frontal gyrus enhances working memory performance

**DOI:** 10.1101/302588

**Authors:** Sankaraleengam Alagapan, Caroline Lustenberger, Eldad Hadar, Hae Won Shin, Flavio Fröhlich

## Abstract

The neural substrates of working memory are spread across prefrontal, parietal and cingulate cortices and are thought to be coordinated through low frequency cortical oscillations in the theta (3 – 8 Hz) and alpha (8 – 12 Hz) frequency bands. While the functional role of many subregions have been elucidated using neuroimaging studies, the role of superior frontal gyrus (SFG) is not yet clear. Here, we combined electrocorticography and direct cortical stimulation in three patients implanted with subdural electrodes to assess if superior frontal gyrus is indeed involved in working memory. We found left SFG exhibited task-related modulation of oscillations in the theta and alpha frequency bands specifically during the encoding epoch. Stimulation at the frequency matched to the endogenous oscillations resulted in reduced reaction times in all three participants. Our results support the causal role of SFG in working memory and suggest that SFG may coordinate working memory through low-frequency oscillations thus bolstering the feasibility of targeting oscillations for restoring cognitive function.

## Introduction

Working memory (WM), the ability to flexibly maintain and manipulate information for a short period of time, forms an important component of cognition. It supports other higher-order cognitive functions and has been tightly linked to fluid intelligence [1, 2]. Impairment in WM accompanies many neurological and psychiatric disorders and significantly reduces the quality of life of affected patients [3-7]. A mechanistic understanding of the causal role of circuit dynamics in WM will open new therapeutic avenues.

Functional imaging studies have revealed that the neural substrate of WM is spread across multiple cortical regions including dorsolateral prefrontal cortex, posterior parietal cortex and anterior cingulate cortex. While early studies have suggested superior frontal gyrus (SFG) to be involved in working memory [8-10], subsequent studies have often found the middle frontal gyrus (MFG) to be the key node in working memory [11-15]. However, lesions in SFG have been shown to result in working memory deficits [16]. In addition, electroencephalography (EEG) and magnetoencephalography (MEG) studies have shown that oscillations in the theta frequency band (4 – 8 Hz) observed on fronto-central regions [17-21] coordinate working memory. The source of these oscillations is thought to be medial prefrontal cortex which includes SFG. Modulations in WM performance by non-invasive brain stimulation like repetitive transcranial magnetic stimulation (rTMS) [22, 23] and transcranial alternating current stimulation (tACS) [24-27] targeting prefrontal cortex also provide indirect evidence for the role of SFG in WM performance.

Electrocorticography (ECoG) allows identification of activity signatures at temporal scale of a few milliseconds with a spatial resolution of a few centimeters is an ideal tool to map functions of cortical regions. Direct cortical stimulation, in which stimulation is applied through ECoG electrodes, allows for focal probing of cortex providing additional information through reversible microlesions [28]. Combined recording and stimulation with implanted electrodes have greatly contributed to revealing the substrate of long-term memory [29-32]. Low amplitude periodic stimulation at 10 Hz has been demonstrated to engage ongoing cortical oscillations in a state-dependent manner and enhance oscillation strength measured by signal power [33]. In this study, we employed a similar experimental paradigm to delineate the role of SFG on working memory. We present results from three participants with subdural electrodes over left and right SFG in whom we assessed the electrophysiological signatures of SFG and applied periodic stimulation during a verbal working memory task. We found that left SFG exhibited a task-related modulation in oscillation power and stimulation matched to the frequency of oscillation resulted in an improvement in working memory performance.

## Results

We leveraged the access to ECoG signals in three patients with epilepsy undergoing long term monitoring in the Epilepsy monitoring unit at the N.C. Neurosciences Hospital, UNC Medical Center, Chapel Hill. The participants (P1, P2 and P3) had electrodes over frontal, temporal and parietal regions on both hemispheres (Figure 1A). The participants performed a Sternberg verbal working memory task that has been previously used in ECoG research [34, 35] (Figure 1B). The cognitive load, measured by the number of items (English letters) in a list to be held in memory (3, 4, or 5 for P1 and 3 or 5 for P2 and 5 or 7 for P3), was varied randomly for each trial. In participant P1, we observed an increase in reaction times with increasing cognitive load (list length 3: 824 ± 31 ms, list length 4: 1119 ± 105 ms, list length 5: 1140 ± 78 ms) in the sham trials (Linear model factor list length: F(2,34) = 4.864; p = 0.014). In participant P2, who performed a separate baseline session of the task without stimulation, there was no significant difference between reaction times for different cognitive loads (F(1,20) = 0.060; p = 0.809). In participant P3, who also performed a separate baseline session without stimulation, there was a significant effect of cognitive load (Linear mixed model factor list length: F(1,45) = 4.646; p = 0.036). The reaction time for trials with 5 items in the list was lower than trials with 7 items in the list (785 ± 26 ms vs 902 ± 48 ms).

**Figure 1.**
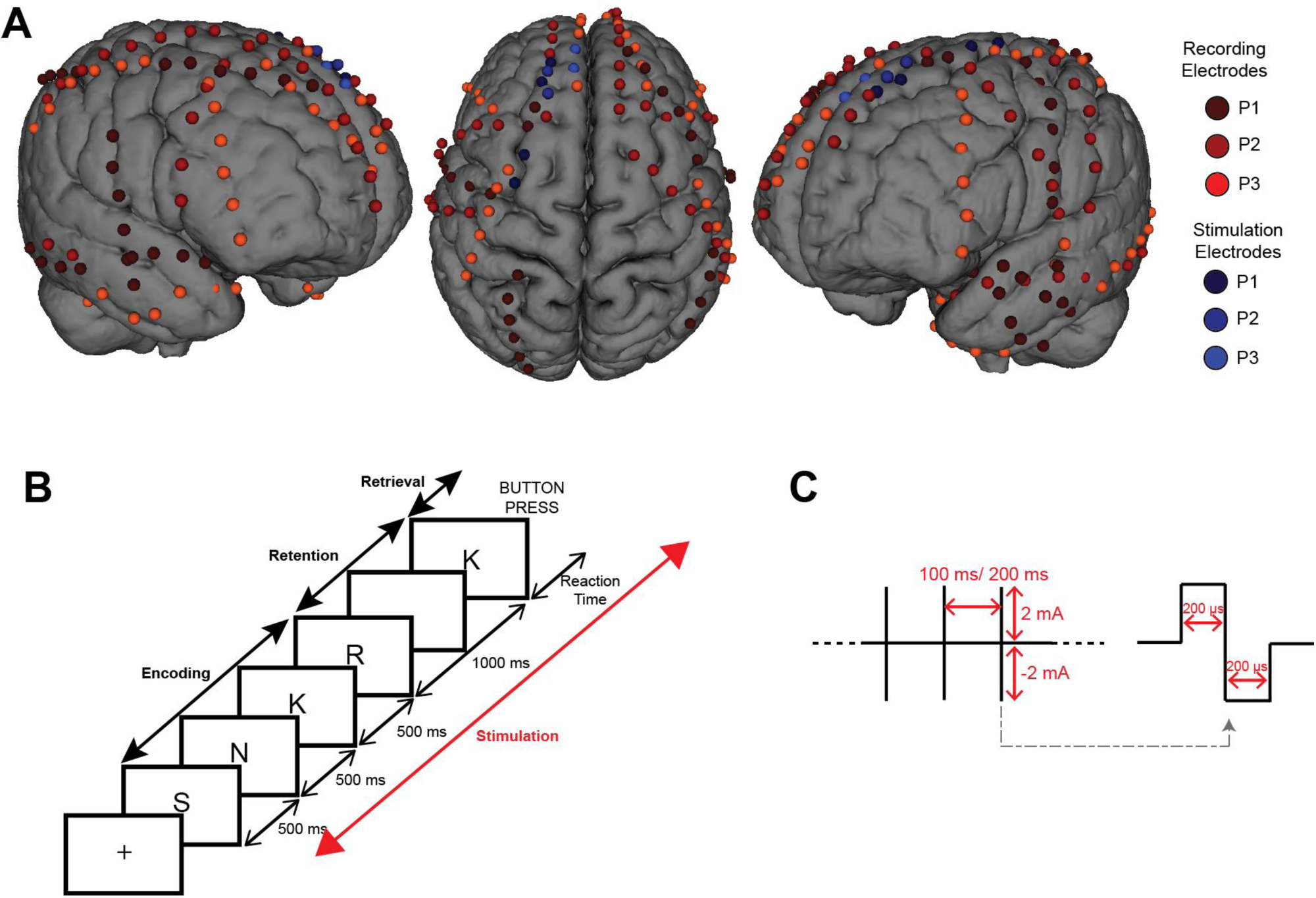
(A) Surface model showing the coverage of electrodes for the three participants. (B) Schematic of a single trial of the working memory task used. The task consisted of 3 epochs – Encoding, Retention and Retrieval. Stimulation was applied through the entire trial. (C) Schematic of the periodic pulse stimulation. Stimulation consisted of train of biphasic pulses 400 μs in duration every 100 ms (P1 and P2) or 200 ms (P3) for 5s.

Oscillations in the theta (3 – 8 Hz) and alpha (8 – 12 Hz) bands have been shown to be modulated during working memory tasks [19, 34, 36]. Spectral analysis revealed oscillations with a peak frequency around 5 Hz in P1 and P3 and 9.5 Hz in P2. To assess if these observed oscillations were modulated by the task, we computed power spectra for baseline, encoding and retention epochs when no stimulation was being delivered (sham trials in P1 and baseline session trials in P2 and P3). Modulation indices were computed relative to baseline epoch. The retrieval epoch was not included in analysis as the epoch may be confounded with action planning and action. We found that electrodes in frontal, temporal and parietal regions exhibited an enhancement of power relative to baseline during sham trials in the theta band (3 – 8 Hz) in P1 and P3 and in alpha band (8 – 10 Hz) in P2 (one sample t-test with FDR correction; p < 0.05). Specifically, electrodes over the left superior frontal gyrus (lSFG) exhibited the task relevant enhancement of oscillation across all three participants. Spectrograms of sample electrodes over SFG illustrating task-related modulation are depicted in Figure 2A. Further analysis of data from electrodes over lSFG revealed that power modulation during the encoding epoch was influenced by list length (Figure 2B; Linear mixed model factor list length: F(3,370) = 3.417; p = 0.017). Post-hoc analysis revealed a significant difference between modulation indices from list lengths 3 and 5 in participants P1 and P2 (Pairwise t-test, p < 0.05). In contrast, power modulation during the retention epoch was not influence by list length (Linear mixed model factor list length: F(3,228) = 1.029; p = 0.38). Taken together, these results imply that the oscillations indeed reflect task relevant processing and specifically contribute to encoding.

**Figure 2.**
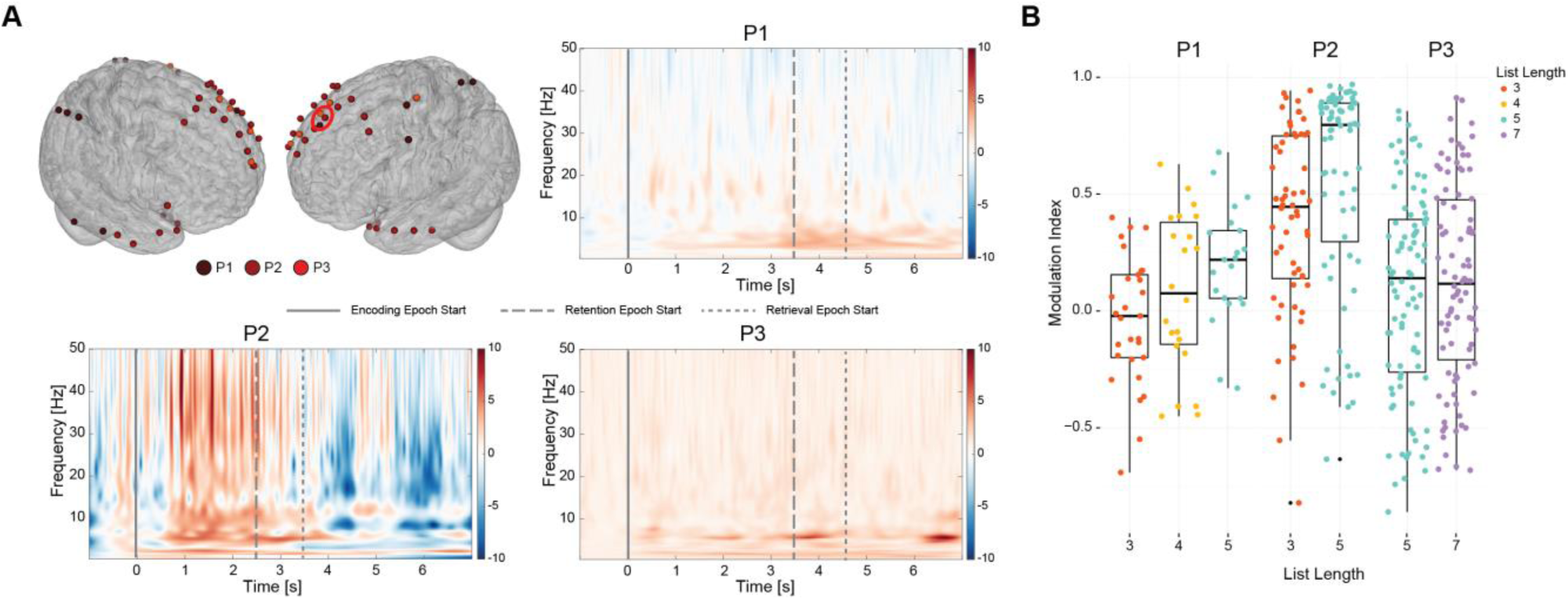
(A) Cortical model showing electrodes that exhibited task-related modulation. Red circle denotes the three electrodes in lSFG whose event related spectral perturbation are plotted. observed in left superior frontal gyrus electrodes during sham trials for P1 and baseline session trials for P2 and P3 indicating the modulation of signal in the band 3 – 12 Hz. Hot (red) colors indicate an increase and cold (blue) colors indicate a decrease in signal power relative to baseline. (B) Modulation indices during encoding epoch across all lSFG electrodes that exhibited significant task related modulation of signal power. In P1 and P2 there was a significant difference between modulation indices for list length 3 and list length 5.

To test if targeting oscillations that exhibited task-related modulation in SFG, periodic pulse stimulation was applied between pairs of electrodes over the left SFG. Stimulation consisted of pulse trains 2 mA in amplitude and 5 seconds in duration (Figure 1C) and the electrode pair being stimulated was randomly changed for each trial. Stimulation and sham trials were randomly interleaved and trial initiation was time-locked to stimulation initiation. Stimulation frequency was 10 Hz for P1 (chosen a priori), 9 Hz for P2 (due to technical issues) and 5 Hz for P3. A total of two different pairs of electrodes in left SFG were stimulated in P1, one pair of electrodes in P2 and P3 (blue electrodes in Figure 1A). During the stimulation session, participants P1, P2 and P3 performed the task in which trials consisted of 3, 4 or 5 items, 3 or 5 items, 5 items respectively. As a first step, the effect of stimulation on reaction times of participants P1 and P2 was analyzed using a linear mixed model with fixed factors list length and stimulation condition and participant as random factor as these participants had trials with 3 and 5 items. While there was no significant effect of stimulation (F(2,108) = 1.042; p = 0.356), there was a significant effect of list length (F(1,108) = 9.072; p = 0.003) and interaction between list length and stimulation condition (F(2,108) = 6.536; p = 0.002). Next analysis was restricted to only trials with 5 items and the reaction times of all 3 participants in sham trials were compared with that in stimulation trials. The effect of stimulation was statistically significant (Linear Mixed Model F(1,97) = 13.414; p <0.001) with all participants showing a significant decrease in reaction times (P1: 1140 ± 78 ms vs 852 ± 111 ms; P2: 1188 ± 93 ms vs 954 ± 54 ms; P3: 841 ± 48 ms vs 727 ± 27ms; Figure 3A) confirmed by post-hoc analysis (Pairwise t-test, p < 0.05). Analysis of accuracy using chi-squared tests did not reveal any significant interactions (Figure 3B) suggesting stimulation served to reduce reaction times without affecting accuracy.

**Figure 3.**
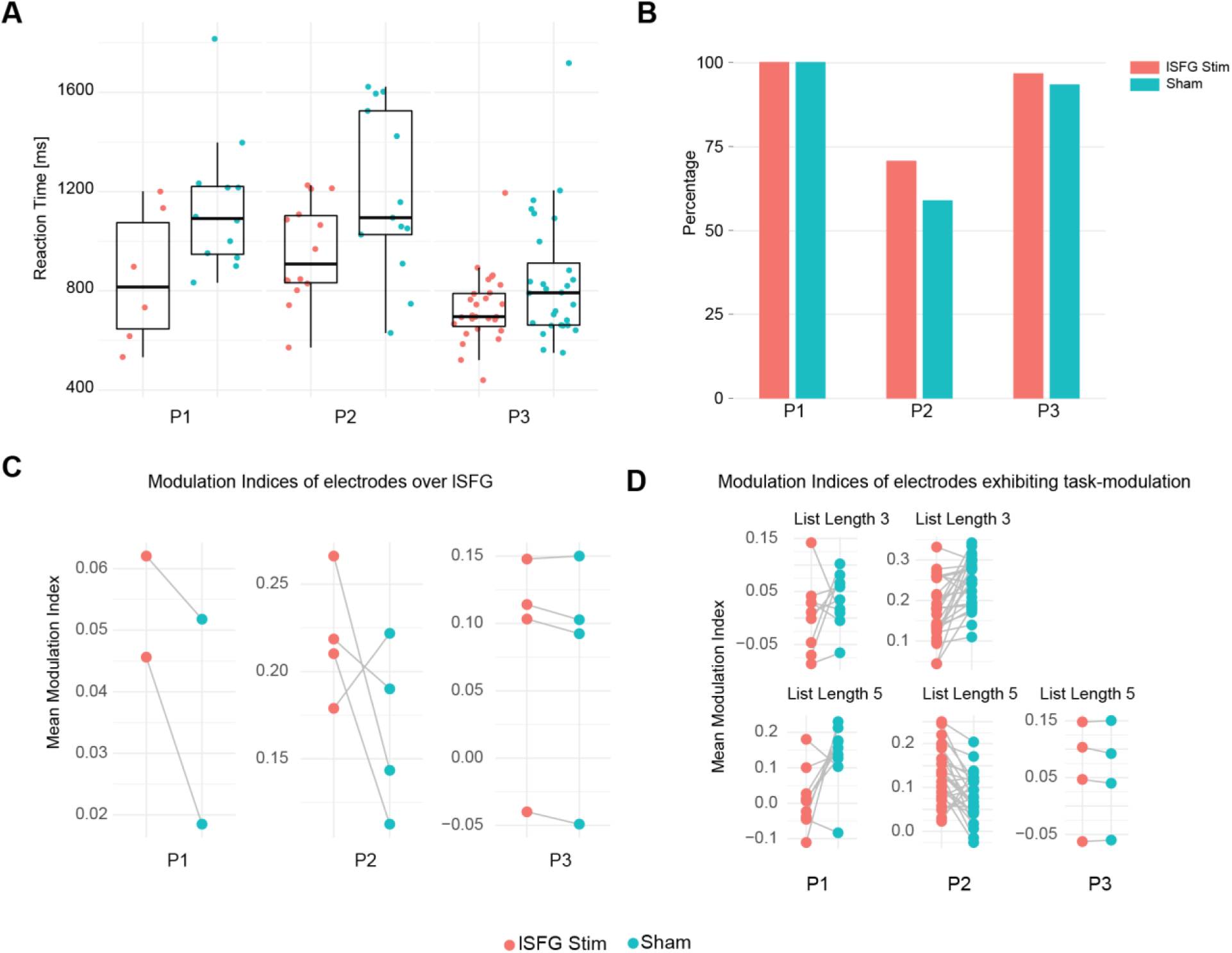
(A) Reaction times in trials with 5 items showing a decrease with stimulation. (B) Accuracy was not affected by stimulation (C) Stimulation did not result in any changes in modulation indices in electrodes over lSFG. (D) Differential effect of stimulation on modulation indices in electrodes that exhibited task-relevant modulation of low frequency oscillations.

In most studies involving electrical stimulation, artifacts caused by stimulation prevent the analysis of electrophysiological signals during stimulation. To overcome this, we developed an independent component analysis (ICA) based method (see Methods and Experimental Procedures). Stimulation artifacts were sufficiently suppressed (Figure S1) allowing us to study the signals in the frequency band of interest. Power spectra and modulation indices in the endogenous oscillation frequency band (3 – 8 Hz in P1 and P3 and 8 – 12 Hz in P2) were computed as described before. Analysis of modulation indices of the electrodes over lSFG (restricted to trials with 5 items in the list) across all participants did not reveal any significant effect of stimulation (Figure 3C; Linear mixed model factor condition F(1,459) = 0.612; p = 0.434). To explore the effects of stimulation on other regions that exhibited modulation of task-relevant oscillations, we ran analysis on individual participant data including list length as a factor. In P1, stimulation induced a differential change in modulation indices (Linear mixed model factor condition F(1,672) = 20.827; p <0.001, factor list length F(1, 672) = 15.793; p = 0.001, interaction F(1,672) = 10.536; p = 0.004). Further analysis revealed that there was a significant effect of stimulation in trials with 5 items in list, with stimulation inducing a decrease in modulation indices (Linear mixed model factor condition F(1,305 = 27.742; p< 0.001). Similarly in P2, stimulation induced a difference change in modulation indices (Linear mixed model factor condition F(1,1738) = 0.495; p = 0.482, factor list length F(1, 1738) = 33.190; p < 0.001, interaction F(1,1738) = 11.134; p < 0.001). Stimulation caused significant decrease in modulation indices in trials with 3 items (Factor condition F(1,908) = 9.04; p = 0.003) while stimulation caused a trend-level significant increase in modulation indices in trials with 5 items (Factor condition F(1,830) = 3.13; p = 0.077). There was no significant effect of stimulation in P3 (Factor condition F(1,215) = 0.005; p = 0.946).

## Discussion

In this study, we show evidence for the role of superior frontal gyrus (SFG) in working memory using a combination of ECoG and DCS. Electrodes over left SFG exhibited modulation of cortical oscillations in the canonical theta and alpha frequency bands. The degree of modulation, measured using modulation index, depended on the cognitive load, specifically in the encoding epoch. Stimulation of lSFG with frequency matched to the fundamental frequency or harmonic of the endogenous oscillations, led to an enhancement in working memory performance. However, analysis of data obtained during stimulation did not provide any conclusive evidence for modulation of task-relevant oscillations. Taken together, the results suggest SFG may be an important node in brain network that coordinates working memory.

While there is an abundance of evidence for the role of middle frontal gyrus (MFG; Brodmann Area 9/46) in working memory from neuroimaging studies [11, 15, 37, 38], the role of SFG is not clear. There have been a few neuroimaging studies that suggest SFG may be involved in working memory [8-10, 39]. SFG gray matter volume has been linked to working memory activation in intra-parietal sulcus [40]. The strongest evidence for the role of SFG in working memory has come from a lesion study [16] in which patients with lesions in lSFG exhibited deficits in working memory involving verbal, spatial and face stimuli. Our results strengthen the evidence for SFG’s role in working memory. However, the proximal location our stimulation targets to MFG may confound our interpretation of the results. Diffusion tensor tractography has revealed that SFG can be divided into subregions with strong connectivity to ACC, a key node in cognitive control network and MFG, a key node in executive control network [41]. As both networks are essential to working memory processes [42-44], stimulation of SFG may have distributed effects across multiple regions including MFG. The lack of sufficient coverage of these areas in these three patients limited our ability to examine this idea. Previous studies have observed oscillations in the range 3 – 15 Hz to be modulated during working memory tasks [34, 45, 46] and the strength of oscillations to reflect working memory load [19, 35, 36]. Frontal midline theta (FMT) is a commonly observed oscillatory signature in EEG studies of working memory [18] typically in Fz and neighboring electrodes in the 10-20 electrode system. The sources of FMT are thought to include lateral PFC and ACC [47]. The theta oscillations we observed in our study may be related to FMT although we did not have any scalp electrodes to confirm this. We found task-related modulation specifically in the encoding period. Analysis of oscillation strength in the retention epoch did not reveal any significant difference between the cognitive loads. This suggests that SFG may play a role that is different from that of MFG/IFG which is known to predominantly be active during the retention epoch [15].

To the best of our knowledge, this is the first study where effects of intracranial stimulation on working memory and on oscillation strength were investigated. Periodic pulse stimulation of entorhinal region has been shown to improve performance in a spatial learning task [31]. Concurrently there was an increase in theta-phase resetting. In another study, stimulation with very weak sinusoidal currents (0.01mA) produced trend level effects in memory performance although no improvement compared to sham was seen [48]. Impairment of performance has been more commonly reported than improvement especially for hippocampal stimulation. One study showed that single pulse stimulation of hippocampus impaired episodic memory [49]. In another study, stimulation at 50 Hz impaired recognition of specific stimuli depending on whether left or right hippocampus was stimulated [50]. More recently, stimulation of entorhinal/ hippocampal and medial temporal regions was shown to affect both verbal and spatial memory [32, 51]. One key difference between the studies described above and our current study is the frequency of stimulation used. Often, 50 Hz was chosen as the stimulation frequency as opposed to the low frequency used in our study. A study that utilized low frequency stimulation showed that stimulation at 5 Hz resulted in improvement of delayed recall [52]. Another study in which theta burst stimulation (100 ms trains of 0.1 ms pulses at 200 Hz repeated 5 times per second) of fornix resulted in improvement of visual-spatial memory [53]. These results suggest that frequency of stimulation might be crucial to the effects observed. Intracranial stimulation studies have often focused on episodic memory and stimulation of hippocampus. In contrast, non-invasive stimulation studies have focused on working memory specifically and target cortical regions such as dlPFC, PPC, inferior frontal gyrus. Transcranial magnetic stimulation, which produces local suprathreshold effects, i.e., evoking action potentials like those expected in intracranial stimulation, has been shown to enhance working memory performance based on the stimulation frequency, location and specific epoch within the task or before the task [54-61]. It must also be noted that many studies report impairments of working memory and episodic memory by TMS as well [62-65]. Transcranial alternating current stimulation, which likely produces more global subthreshold effects, has been shown to increase performance by targeting dlPFC and PPC [24, 26]. The neurophysiological underpinnings of the effects in these studies are often unclear [27, 60]. Recently, rTMS applied at theta frequency to left intraparietal sulcus was shown to entrain theta oscillations with a concurrent improvement in auditory working memory [66].

As any scientific study, our study has a set of limitations. First, the results presented here are from three participants. The major obstacle in our case was the heterogeneity in electrode distribution as the electrode locations were dictated by clinical needs. Second, although the stimulation frequency was 10 Hz, oscillations in the frequency band 3 – 8 Hz were significantly modulated concurrently with changes in WM performance. This discrepancy is hard to reconcile if entrainment is thought to be the underlying mechanism of interaction between stimulation and oscillation [67, 68]. However, the interaction between stimulation and an ongoing oscillation has been found to be nonlinear and the effects depend on the strength of the prevailing oscillations [33]. When there is a strong ongoing oscillation, stimulation tends to increase the strength of the endogenous oscillation and only in cases where the strength of the oscillation is low, entrainment is possible. This state-dependent effect of stimulation is likely the underlying mechanism in the current study as well. Alternatively, 10 Hz stimulation may have engaged with the strong 5 Hz oscillation through subharmonic entrainment as predicted in computational models [69].Third, the present experimental paradigm is limited to applying stimulation during the entire trial due to technical limitations of the FDA-approved cortical stimulator used in the study. This limitation precluded us from identifying if stimulation during an epoch within a trial, i.e. encoding or retention, is more effective than stimulation during the entire trial. Moreover, the frequency of stimulation was restricted to a few discrete frequencies that did not allow matching of the stimulation frequency to frequency of endogenous oscillations in P1. Fourth, a limitation of the current study design is that it used only a single stimulation amplitude and stimulation frequency. Given the large parameter space, it is prohibitively difficult to try all possible parameters in studies with limited participant pools as the current study. For P1, we chose stimulation regions based on previous literature due to technical limitations. A more effective strategy was followed for P2 and P3 where we identified electrodes that exhibited task-related modulation in low frequency bands and applied stimulation accordingly. Also, the stimulation used in our study was restricted to a single site. However, memory processes are distributed across different brain regions and the most effective strategy would likely involve stimulation of multiple regions to produce more of a network effect [29, 70] or an adaptive approach using closed-loop stimulation based on the state of the network [71, 72]

In conclusion, we show that periodic pulse stimulation of cortex through subdural electrodes at low frequency can enhance working memory. Despite the limitations, the study provides valuable insights into the feasibility of using oscillations as brain stimulation targets. The importance is highlighted by the emerging interest in using invasive recordings and electrical stimulation to understand and alter pathological signatures of brain activity, whether it be neurological disorders, like epilepsy and Parkinson’s disease, or psychiatric disorders, like depression and obsessive-compulsive disorder. Our results suggest that the same technology could be leveraged to also address cognitive impairment.

### Experimental Procedures

#### ECoG Data Collection and Direct Cortical Stimulation

All experimental procedures were approved by the Institutional Review Board of University of North Carolina at Chapel Hill (IRB Number 13-2710) and written informed consent was obtained from the participant. The participants underwent implantation of intracranial EEG electrodes followed by long-term monitoring at the Epilepsy Monitoring Unit in UNC Neuroscience hospital for surgical resection planning.

Strips of electrodes were implanted over bilateral frontal, temporal and parietal lobes as shown in Figure 1A. Depth electrodes were implanted in bilateral parahippocampal gyri in P1 and strip electrodes were implanted over bilateral occipital lobe in P2 (not shown in figure). The locations of the electrodes were completely dictated by the clinical needs of the participant. The electrodes, 4 mm in diameter (2.5 mm exposed), were made of platinum-iridium alloy and embedded in silicone (Ad-Tech Medical, Racine, Wisconsin, United States). The electrodes in each strip were separated by 10 mm. Signals from electrodes that were over seizure foci (Table 1) were excluded from analysis.

**Table 1.** Clinical Information of Participants

| Participant | Age | Sex | Handedness | Clinical Seizure Focus | Stimulation Frequency | Number of Trials |
| --- | --- | --- | --- | --- | --- | --- |
| P1 | 23 | F | R | Bilateral parahippocampal gyri | 10 Hz | 24 Sham, 13 Stimulation |
| P2 | 57 | M | R | Bilateral inferior occipital, posterior temporal | 9 Hz | 27 Sham, 26 Stimulation |
| P3 | 26 | M | R | Unknown Seizure Focus | 5 Hz | 30 Sham, 30 Stimulation |

ECoG data from participant P1 was recorded using a 128-channel acquisition system (Aura LTM 64, Grass Technologies, Warwick, Rhode Island, United States) at 800 Hz sampling rate. Electrical stimulation consisted of 5 second train of biphasic pulses, 2 mA in amplitude, 400 μs in duration and 10 Hz in frequency. The pulses were generated by a cortical stimulator (S12x cortical stimulator, Grass Technologies, Warwick, Rhode Island, United States) and applied between pairs of adjacent electrodes (blue electrodes in Figure 1A).

ECoG data from participants P2 and P3 were recorded using a different 128-channel EEG system (NetAmps 410, Electrical Geodesics Inc, Eugene, Oregon, United States) at 1000 Hz sampling rate. Stimulation was delivered using Cerestim M96 cortical stimulator (Blackrock Microsystems, Salt Lake City, Utah, United States). Stimulation parameters (except frequency) remained the same as in P1 except for the duration which was adjusted to encompass the encoding and retention epochs.

#### Working Memory Task

We adopted a classical Sternberg working memory task previously used in other ECoG studies [34, 35, 73] (Figure 1C). The task consisted of 3 epochs. In the first epoch, lists of 3 to 5 pseudo-randomly chosen letters from the English alphabet were presented sequentially. This was termed the encoding epoch and each alphabet was displayed for 500 ms with 200 ms between each alphabet (the inter-alphabet interval was not present for P2 and P3). Following this, was a retention epoch where a blank screen was presented for 1 second. The final epoch was the retrieval epoch where a single probe (another English alphabet) was shown for 5 seconds and the participants had to indicate if they thought that the probe was present in the list by pressing a specified key on the keyboard. If they did not think the probe was present in the list, they did not have to press any key. The task was programmed in Matlab using Psychtoolbox [74] and presented in a laptop. For the experiment in which P1 participated, triggers from the cortical stimulator were detected by an ethernet DAQ (National instruments, Austin, TX, USA) connected to the task computer and used to initiate trials. Sham trials, in which no electrical pulses were delivered, were initiated using a pulse generator and were randomly interleaved with stimulation trials. For the experimental session in which P2 and P3 participated, triggers were generated within the Psychtoolbox task code and sent to Cerestim through the ethernet DAQ. In sham trials, no triggers were sent to Cerestim. Stimulation was applied for 5 seconds in P1 and the duration of encoding and retention epochs in P2 and P3. In P1 electrodes over right SFG and bilateral temporal cortices were stimulated as well. However, the low number of stimulation trials did not allow any meaningful analysis to be performed and hence was not included in the study here. In P2, a pair of electrode over right SFG was stimulated and the results are not included here.

Participants P2 and P3 completed 2 sessions – a baseline session and a stimulation session. The baseline session did not include any stimulation and consisted of 40 trials of two different list lengths to assess the baseline performance level as well as determine the parameters for the stimulation session.

#### Data Analysis

Data analysis was performed using custom written Matlab scripts (The MathWorks Inc., Natick, MA, United States). The recording setup consisted of switching circuits designed to protect the amplifier during stimulation which prevented recording of data from stimulating electrodes. Hence, data from stimulating electrodes were not included in the analysis.

Stimulation artifacts, present in channels adjacent to stimulated electrodes, were removed using an independent component analysis (ICA) based approach (Figure S). Since artifacts were observed as stereotypical waveforms, ICA resulted in components that contained only artifact waveforms which were then rejected, and the remaining components were used to reconstruct artifact free signals. We used the infomax algorithm [75] available as a part of EEGLab toolbox [76] for computing independent components. Following artifact suppression, the signals were low pass filtered with an FIR filter (cutoff frequency 50 Hz) and re-referenced to common average. Signal power spectra was computed with a multi-taper fft based approach using Chronux toolbox [77]. To quantify the change induced by stimulation, modulation index was computed as

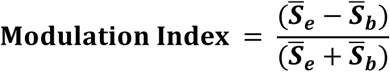

Where ***S_e_*** and ***S_b_*** are average power in specified frequency band in specific epoch (task, encoding or retention) and baseline epoch respectively. The baseline epoch was defined as 5 second interval before the beginning of encoding epoch.

Time-frequency representations were computed by convolving Morlet wavelets with the time series of each trial. Event related spectral perturbation was calculated as

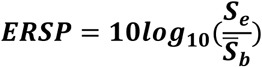

Where ***S_e_*** is the spectra at each time point within an epoch and ***S_b_*** is the average power in the baseline epoch.

#### Statistics

All statistical analyses were performed using R. Linear mixed effects models were fitted using the lmertest package [78] which uses Satterthwaite’s approximation to degrees of freedom to determine the F statistics of the fixed effects.

For the effect of list length on reaction times, we fitted a linear model with reaction time as dependent variable and list length as the factor for each participant separately. For the effect of list length on modulation indices, we fitted linear mixed model with modulation index as the dependent variable and list length as the fixed factor and participant and electrodes as nested random factors. To study the effect of stimulation on reaction time, we fitted a linear mixed model with reaction time as dependent variable and stimulation as fixed factors and participant as the random factor. As post hoc analysis we performed a two-sample t-test to compare the difference between reaction times during sham and stimulation trials for each participant. To study the effect of stimulation on modulation index, we fitted linear mixed models with modulation index as dependent variable, stimulation as fixed factor and electrodes and participants as nested random factors and also with modulation index as dependent variable and list length (3 levels) and stimulation regions (3 levels – sham, frontal region, temporal region) as fixed factors and electrodes as a random factor. As post-hoc analysis, we performed paired t-tests.

#### Extraction of Electrode Location from Neuroimaging Data

3D Slicer [79] was used to analyze and extract electrode locations from CT images obtained after implantation of subdural electrodes. The post-operative MRI was co-registered to post-operative CT in Slicer followed by registering to standard MNI atlas [80]. Skull stripping was performed using ROBEX [81], and the gray matter and white matter were then segmented using ITK-Snap [82]. The surface model of the MNI atlas brain was generated using Slicer and used for visualization purposes. The anatomical locations of the electrodes were determined by co-registering the MRI Image to the MNI Atlas [83], recomputing electrode locations in the MNI space, transforming these locations to Talairach space, and using the Talairach Client [84] to obtain the label of the gray matter nearest to the coordinate representing electrode location.

## Acknowledgments

The authors thank the members of the Frohlich Lab for their valuable input, with special thanks to Sangtae Ahn for verifying the accuracy of the codes used for analysis and providing valuable feedback on the manuscript. The authors also thank the EEG technicians at the UNC Epilepsy monitoring unit for their generous help with the ECoG recordings.

## Funding

Research reported in this publication was supported in part by the National Institute of Mental Health of the National Institutes of Health under Award Numbers R01MH101547 and R21MH105557, National Institute of Neurological Disorders and Stroke of the National Institutes of Health under award number R21NS094988-01A1, Translational Team Science Award (TTSA) with funding provided by the UNC School of Medicine and the National Center for Advancing Translational Sciences (NCATS), National Institutes of Health, through Grant Award Number UL1TR001111, Helen Lyng White Postdoctoral Fellowship (S.A.) and Swiss National Science Foundation (grant P300PA_164693; C.L.). The content is solely the responsibility of the authors and does not necessarily represent the official views of the National Institutes of Health.

## Supplementary Figures

**Figure S1:**
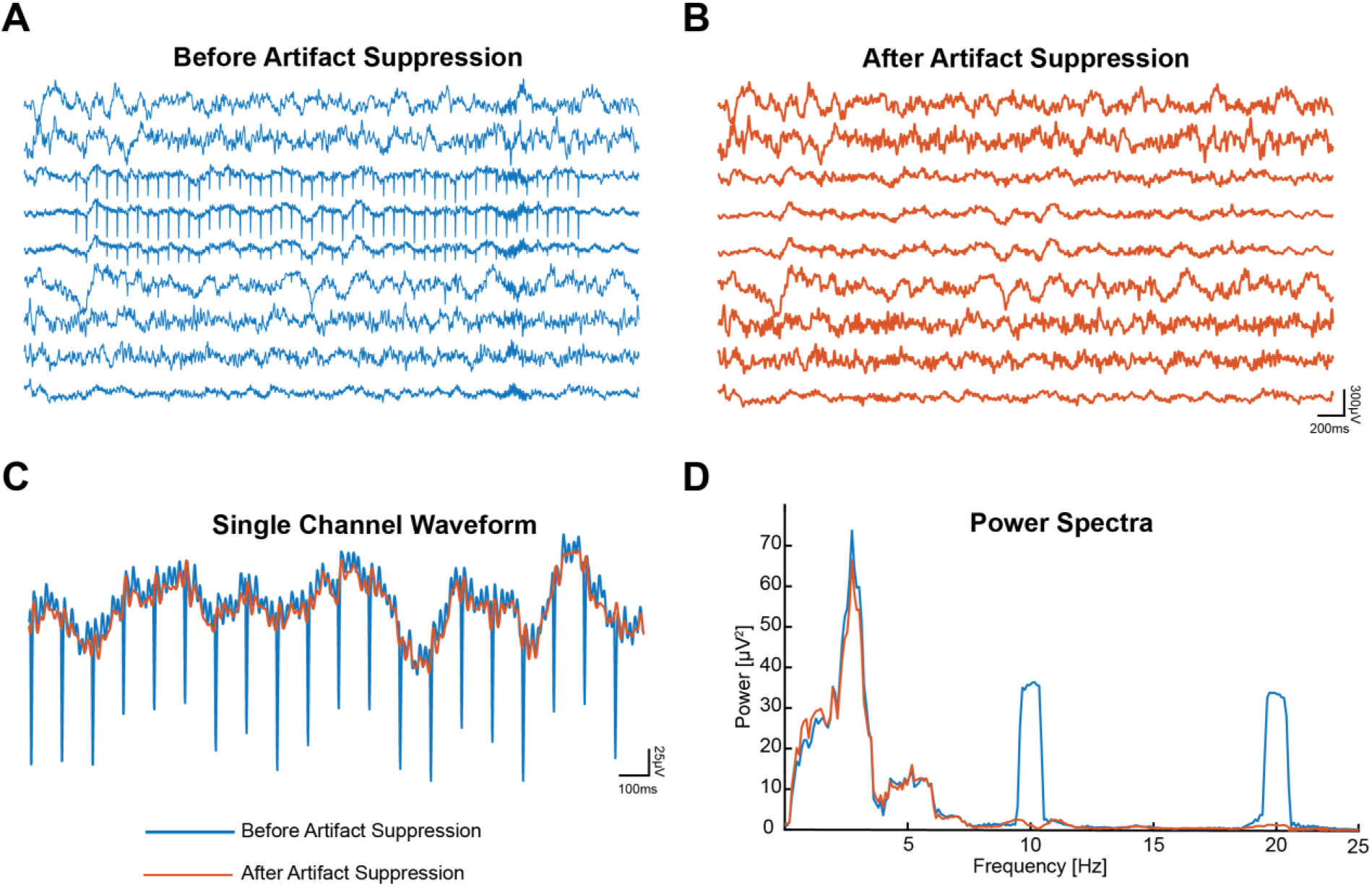
An example illustrating performance of ICA based artifact suppression algorithm. (A) Raw signal from channels adjacent to stimulation channels before artifact suppression (B) Signal after removal of stimulation artifacts. Visually, the artifact has been reduced to noise level. (C) Trace from a single channel highlighting the suppression of artifact waveform (D) Power spectra computed from the signal shown in (C) before and artifact suppression.

